# Functionally critical residues in the aminoglycoside resistance-associated methyltransferase RmtC play distinct roles in 30S substrate recognition

**DOI:** 10.1101/712810

**Authors:** Meisam Nosrati, Debayan Dey, Atousa Mehrani, Sarah E. Strassler, Natalia Zelinskaya, Eric D. Hoffer, Scott M. Stagg, Christine M. Dunham, Graeme L. Conn

## Abstract

Methylation of the small ribosome subunit rRNA in the ribosomal decoding center results in exceptionally high-level aminoglycoside resistance in bacteria. Enzymes that methylate 16S rRNA on N7 of nucleotide G1405 (m^7^G1405) have been identified in both aminoglycoside-producing and clinically drug-resistant pathogenic bacteria. Using a fluorescence polarization 30S-binding assay and a new crystal structure of the methyltransferase RmtC at 3.14 Å resolution, here we report a structure-guided functional study of 30S substrate recognition by the aminoglycoside resistance–associated 16S rRNA (m7G1405) methyltransferases. We found that the binding site for these enzymes in the 30S subunit directly overlaps with that of a second family of aminoglycoside resistance–associated 16S rRNA (m^1^A1408) methyltransferases, suggesting both groups of enzymes may exploit the same conserved rRNA tertiary surface for docking to the 30S. Within RmtC, we defined an N-terminal domain surface, comprising basic residues from both the N1 and N2 subdomains, that directly contributes to 30S-binding affinity. In contrast, additional residues lining a contiguous adjacent surface on the C-terminal domain were critical for 16S rRNA modification, but did not directly contribute to the binding affinity. The results from our experiments define the critical features of m^7^G1405 methyltransferase–substrate recognition and distinguish at least two distinct, functionally critical contributions of the tested enzyme residues: 30S-binding affinity and stabilizing a binding-induced 16S rRNA conformation necessary for G1405 modification. Our study sets the scene for future high-resolution structural studies of the 30S–methyltransferase complex and for potential exploitation of unique aspects of substrate recognition in future therapeutic strategies.

---

Methylation of 16S rRNA has been identified as a prominent mechanism of self-protection in aminoglycoside-producing bacteria and is emerging as a new threat to the clinical efficacy of aminoglycoside antibiotics (1, 2). Both the intrinsic methyltransferases of drug producers and acquired enzymes of human and animal pathogens chemically modify the aminoglycoside binding site in the decoding center of the bacterial 30S subunit to block drug binding and confer exceptionally high-level resistance. Regarding the acquired enzymes specifically, of most concern is that these resistance determinants have been identified on various mobile genetic elements, often in conjunction with other resistance enzymes (2–4). As such, the aminoglycoside-resistance methyltransferases can make the bacteria expressing them pan-resistant to entire subclasses of aminoglycosides (2, 5), including even the most recent generation drugs like plazomicin (6, 7). More broadly, given the extensive modification of bacterial rRNAs, especially in functionally critical regions like the decoding center, understanding rRNA methyltransferase-ribosome subunit interactions has relevance to both fundamental bacterial physiology as well as mechanisms of antimicrobial resistance.

The aminoglycoside-resistance 16S rRNA methyltransferases are functionally divided into two subfamilies that modify the ribosome at either the N7 position of 16S rRNA nucleotide G1405 (m^7^G1405) or the N1 position of A1408 (m^1^A1408). While enzymes from both subfamilies are found in aminoglycoside-producing bacteria, the m^7^G1405 methyltransferases (**Fig. 1A**) are far more clinically prevalent than their m^1^A1408 methyltransferase counterparts (2, 8). In contrast to the single m^1^A1408 methyltransferase NpmA that was clinically isolated from *E. coli* strain ARS in Japan (9), the m^7^G1405 methyltransferases are globally disseminated and have been found in many different human pathogens (2).

**Fig. 1.**
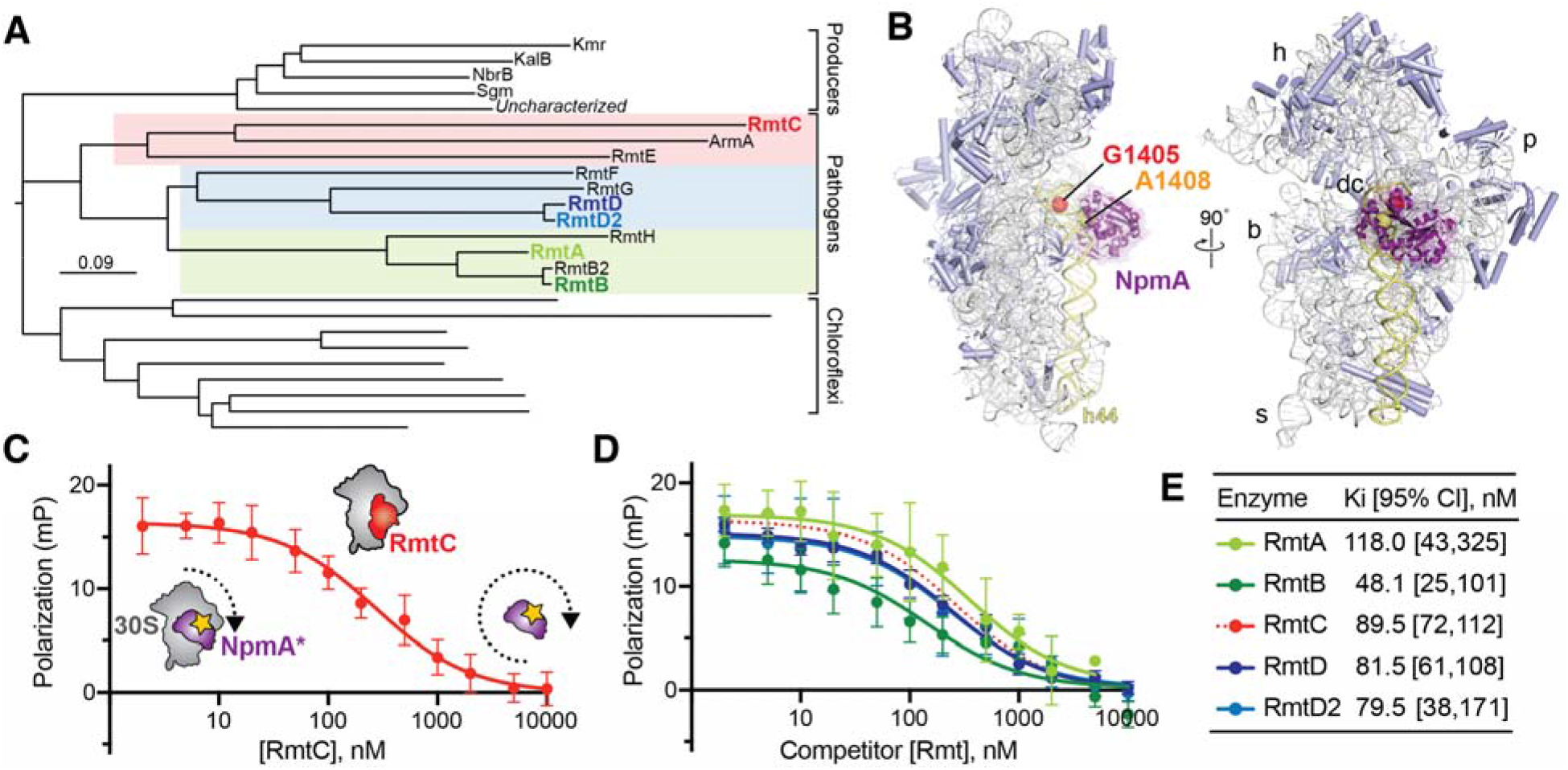
The m^7^G1405 methyltransferase family and binding site on 30S subunit. **A**. Phylogenetic tree of m^7^G1405 methyltransferase enzymes including acquired genes in gammaproteobacterial and Gram-negative pathogenic bacteria, aminoglycoside producing bacteria and uncharacterized homologs belonging to the chloroflexi. Pathogen-associated genes (color-coded regions) are further divided into three subclades represented in this work by RmtA/ RmtB, RmtD/ RmtD2 and RmtC. **B.** Structure of the bacterial 30S subunit bound to NpmA (purple) (PDB ID: 4OX9) showing the proximity of nucleotides G1405 (red) and A1408 (orange) at the top of h44 (yellow) in the ribosome decoding center (dc). Other 30S features indicated are the head (h), body (b), platform (p), and spur (s). **C.** Schematic of the competition FP assay for 30S-methyltransferase binding using the NpmA* probe (purple) and application to RmtC binding (red). At low competitor concentration (left of plot) high FP signal arises due to NpmA* interaction with 30S; displacement of the probe by RmtC results in lower FP signal from the free probe (right of plot). **D, E.** Competition FP binding experiments with NpmA* and five different unlabeled wild-type Rmt enzymes. The RmtC curve in panel D (red dotted line) is the same as in panel C and is shown for comparison. Binding affinities (K_i_) and associated 95% confidence interval were obtained from fits to the data shown in panels C and D; Error bars represent SD of the measurements.

Both free and 30S-bound m^1^A1408 methyltransferases, including NpmA, have been extensively characterized, revealing the molecular basis of their specific substrate recognition and modification mechanisms (10–15). These enzymes exploit a conserved 16S rRNA tertiary surface adjacent to helix 44 (h44) to dock on the 30S, explaining the requirement for intact 30S as their substrate. Two extended regions that connect the fifth/ sixth and sixth/ seventh β-strands of the methyltransferase core fold (β5/6 and β6/7 linkers, respectively) position key residues for recognition and stabilization of A1408 in a flipped conformation for methylation (10, 13).

Structures of the m^7^G1405 methyltransferases RmtB (16), which has been identified in multiple Gram-negative pathogens, and Sgm (17), from the producer of sisomicin derivative G52, *Micromonospora zionensis*, have revealed a distinct methyltransferase architecture. Specifically, these enzymes possess a significantly larger N-terminal extension but no extended sequences within the methyltransferase core fold comparable to those in the m^1^A1408 methyltransferases. A likely role for the unique N-terminal domain in 30S interaction by the m^7^G1405 methyltransferases has been suggested and some functionally critical residues within this domain have been previously identified (16–19). However, to date, no direct binding analysis to allow dissection of important residues in binding or stabilization of a catalytically competent state of the enzyme-substrate complex has been performed. As such, there is a critical gap in our understanding of m^7^G1405 methyltransferases 30S substrate recognition, despite the potential threat these enzymes pose for clinical aminoglycoside resistance.

Here, we have extended the use of a 30S binding assay previously developed in our lab for studies of NpmA (13) to the m^7^G1405 methyltransferases. From these direct 30S binding measurements and a structure-guided mutagenesis strategy based on a new structure of a m^7^G1405 methyltransferase family member (RmtC), we develop a new model for 30S substrate recognition by the m^7^G1405 methyltransferases. We identify a molecular surface in the N-terminal domain that is critical for 30S docking, while numerous residues on an adjacent surface of the CTD do not contribute to binding affinity but likely control critical conformational changes necessary for catalysis of rRNA modification.

## RESULTS

### m^7^G1405 methyltransferases bind 30S with similar affinity and at a site overlapping that of the m^1^A1408 methyltransferases

We previously developed a competition fluorescence polarization (FP) assay to measure the binding affinity of wild-type and variant NpmA proteins to define this methyltransferase’s mechanism of 30S substrate recognition and m^1^A1408 modification (13). We speculated that the close proximity of nucleotides A1408 and G1405 in h44 (**Fig. 1*B***) might also make this assay applicable to direct quantification of m^7^G1405 methyltransferase-30S interactions. In this assay, a fluorescein-labeled, single-Cys variant (E184C) of NpmA (NpmA*) is prebound to 30S (high FP state) and a range of concentrations of unlabeled competitor protein added to displace the NpmA* probe (shown schematically in **Fig. 1*C***), allowing determination of the methyltransferase 30S-binding affinity (K_i_). We first applied the assay to analysis of 30S-RmtC interaction and observed a RmtC-concentration dependent decrease in FP. The resulting data were fit to obtain a K_i_ of 89.5 nM (**Fig. 1*C,E***). This value is comparable to the 60 nM affinity previously measured for the m^1^A1408 methyltransferase NpmA (13). Binding measurements were also performed with RmtA, RmtB, RmtD and RmtD2, which together with RmtC, represent each of the three subclades in the m^7^G1405 methyltransferase phylogenetic tree (**Fig. 1*A***). All binding affinities for these methyltransferases were comparable within a ~2.5-fold range from 48 to 118 nM (**Fig. *D,E***).

These results confirm our established assay using the NpmA-E184C* probe as suitable for direct binding measurements of m^7^G1405 methyltransferases to 30S and thus as a tool to provide a deeper analysis of their substrate recognition mechanism. These data also reveal that the binding site of the m^7^G1405 methyltransferases on the 30S subunit does indeed overlap with that of the m^1^A1408 methyltransferases, suggesting they may also exploit the same conserved rRNA tertiary surface for specific substrate recognition. We chose to use RmtC for further structural and functional studies of 30S-m^7^G1405 methyltransferase interaction for several reasons. Most importantly, there has been no such analysis of RmtC to date and this enzyme is both in the same subclade as ArmA and most distant from RmtB (**Fig. 1*A***), two commonly observed pathogen-acquired m^7^G1405 methyltransferases. The selection of RmtC thus offers the opportunity to identify conserved features of the 30S recognition mechanism across all m^7^G1405 methyltransferases.

### Structure of the RmtC-SAH complex

The X-ray crystal structure of RmtC bound to S-adenosylhomocysteine (SAH), the methylation reaction by-product, was determined and refined at 3.14 Å resolution (**Table 1**). RmtC adopts a fold consistent with those of other m^7^G1405 methyltransferases RmtB and Sgm (16, 17), as expected. Specifically, RmtC possesses a large amino-terminal domain (NTD) appended to its carboxy-terminal domain (CTD) methyltransferase fold (**Fig 2*A,B***). The NTD is structurally divided into two subdomains, N1 and N2, each comprised of three ⍰-helices. N1 forms a globular three-helical bundle, while the three helices of N2 are extended across the N-terminal half of the CTD (**Fig. 2*A***). The CTD adopts a canonical Class I methyltransferase fold with a seven-stranded β-sheet core containing a central topological switch point that forms the SAM binding pocket (**Fig. 2*B***).

**Table 1.** X-ray data collection and structure refinement statistics

|  | RmtC+SAH |
| --- | --- |
| PDB ID | 6PQB |
| Space group | P61 |
| Resolution (Å) | 40.9-3.14 (3.25-3.14)<sup>a</sup> |
| Cell dimensions |  |
| *a*, *b*, *c* (Å) | 163.5, 163.5, 122.5 |
| $\alpha$, $\beta$, $\gamma$ (°) | 90, 90, 120 |
| Molecules a.s.u. | 4 |
| Wavelength, Å | 0.987 |
| *R*<sub>pim</sub> | 0.047 (0.867) |
| *CC*<sub>1/2</sub> | 0.998 (0.419) |
| *I* / $\sigma I$ | 10.81 (1.34) |
| Completeness (%) | 100 (100) |
| Redundancy | 8.7 (8.2) |
| Total reflections | 283,111 |
| Unique reflections | 32,566 |
| *R*<sub>work</sub> / *R*<sub>free</sub><sup>b</sup> | 20.0/23.3 |
| Atoms |  |
| Protein | 8538 |
| SAH | 104 |
| Water | 9 |
| Average B-factor | 134 |
| Ramachandran, % |  |
| Favored/ allowed | 99.8 |
| Disallowed | 0.2 |
| R.m.s. deviations |  |
| Bond lengths (Å) | 0.002 |
| Bond angles (°) | 0.474 |

<sup>a</sup>Values in parenthesis are for the highest resolution shell.
<sup>b</sup> $R_{\text{work}} = \sum hkl |F_o(hkl) - F_c(hkl)| / \sum hkl |F_o(hkl)|$ , where *F*<sub>o</sub> and *F*<sub>c</sub> are observed and calculated structure factors, respectively. *R*<sub>free</sub> applies to the 5% of reflections chosen at random to constitute the test set

**Fig. 2.**
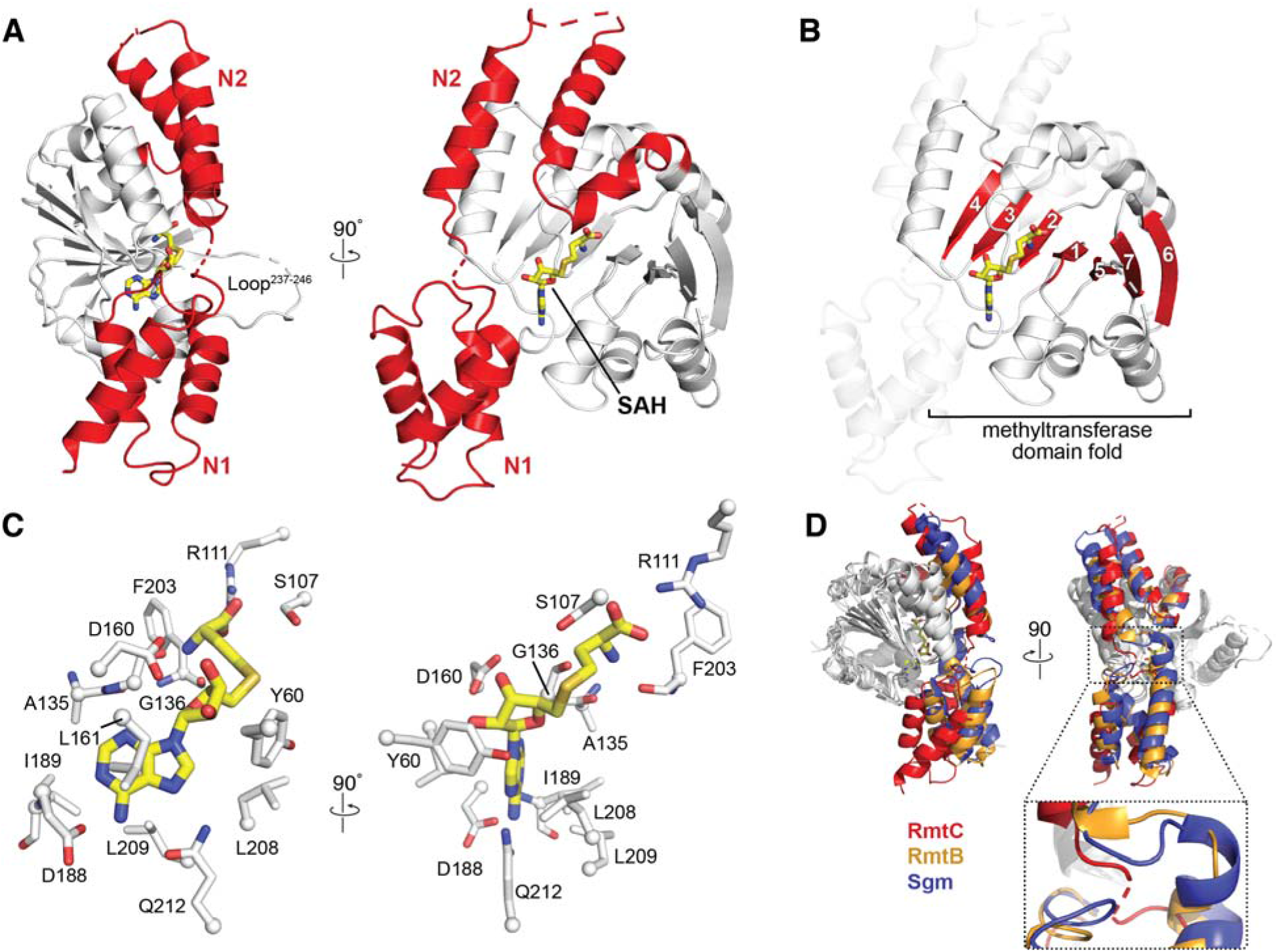
Structure of the RmtC-SAH complex. **A.** Crystal structure of the RmtC-SAH highlighting (red) the extended N-terminal domain characteristic of the aminoglycoside-resistance 16S rRNA (m^7^G1405) methyltransferases. The N-terminal domain is divided into two subdomains, N1 and N2. The locations of the bound SAH (yellow sticks) and a partially disordered loop (Loop^237-246^) adjacent to the opening to the SAM-binding pocket, are also indicated. **B.** The same view of the RmtC structure as panel A (*right*) but highlighting the seven β-strand core (red) of the C-terminal methyltransferase fold (with N1 and N2 shown as semitransparent cartoon). **C.** Two orthogonal detailed views of the interactions made with SAH in the SAM-binding pocket. **D.** Alignment of RmtC (red) with the structures of RmtB (PDB ID: 3FRH; orange) and Sgm (PDB ID: 3LCV; blue), shown in two orthogonal views, *top*, reveal potential flexibility in the position of the N1 domain relative to the N2/CTD domains via a hinge region between N1 and N2 (zoomed view).

In the RmtC-SAH complex, the SAH is bound in a pocket lined by numerous conserved residues, with numerous hydrogen bonding and hydrophobic interactions. Two highly conserved residues, Arg111 and Asp160, anchor the SAH carboxylate group and ribose hydroxyl groups, respectively (**Fig. 2*C***), while Asp188 and Gln212 position the base via hydrogen bonds to the adenine amino group and ring N7. The SAH ribose and adenine moieties are also surrounded by a collection of hydrophobic side chains on each side that define the shape of the binding pocket. Overall, the interactions made by RmtC with SAH in the SAM binding pocket are consistent with previous structures of RmtB and Sgm bound to cosubstrate (16, 17). During the course of this work, a structure of *apo* RmtC (PDB ID: 6CN0) was also deposited by the Center for Structural Genomics of Infectious Diseases. Comparison of the RmtC-SAH complex with this structure reveals the interactions within the SAM binding pocket to be mostly maintained. However, some potential conformational flexibility is apparent in residues Tyr60 and Ser107. These residues line the opening to the SAM binding pocket and may assist in positioning G1405 close to the SAM methyl group for modification (17).

Structural alignment of our RmtC structure with those of RmtB and Sgm confirms these proteins are structurally similar overall (average RMSDs of 2.62 and 3.05 Å, respectively). However, a substantial difference in the orientation of the N1 subdomain relative to the remainder of the protein is apparent in alignments made using only the CTD of each structure (**Fig. 2*D***), reducing the average RMSDs to 1.59 and 1.50 Å, respectively. Additionally, at least two residues in all four copies of RmtC in the crystal are disordered in the sequence that links N1 and N2 (between positions 62 and 64). Together, these observations suggest the potential for flexibility in N1 subdomain position relative to the remainder of the protein and the sequence between N1 and N2 may act as a hinge that allows movement of this subdomain (**Fig. 2*D***). Given the essential role of the N1 domain in substrate binding (see below), such mobility between the N1 and N2 domains may be an important aspect of specific 30S substrate recognition.

### Identification of potential 16S rRNA-binding residues in RmtC

Previous studies of Sgm, ArmA and RmtB identified the importance of the m^7^G1405 methyltransferase NTD in substrate recognition and have also suggested a specific role in 30S binding for some residues within both protein domains (16–19). The likely importance of conserved positive surface charges in the NTD are further supported by our structure of RmtC in which residues of the N1 and N2 subdomains form an extended, contiguous positively charged surface that could interact with 16S rRNA (**Fig. 3*A***). Previous structure-guided mutagenesis of RmtB coupled with tobramycin minimum inhibitory concentration (MIC) assays identified several residues potentially important for 30S binding (16), including highly conserved residues within a structurally disordered loop (corresponding to RmtC residues 237-246; Loop^237-246^). In the SAH-bound structure of RmtC, like the previously determined structures of RmtB and Sgm (16, 17), there is weak or no density visible for most Loop^237-246^ residues, including the highly conserved Lys236 and Arg241. The functional importance of these and other conserved residues in the absence of an obvious role in Rmt protein structure or SAM binding is suggestive of an important contribution to 30S substrate recognition. However, to date, no measurements of 30S binding have been made for any m^7^G1405 methyltransferase to directly test the roles of these important residues.

**Fig. 3.**
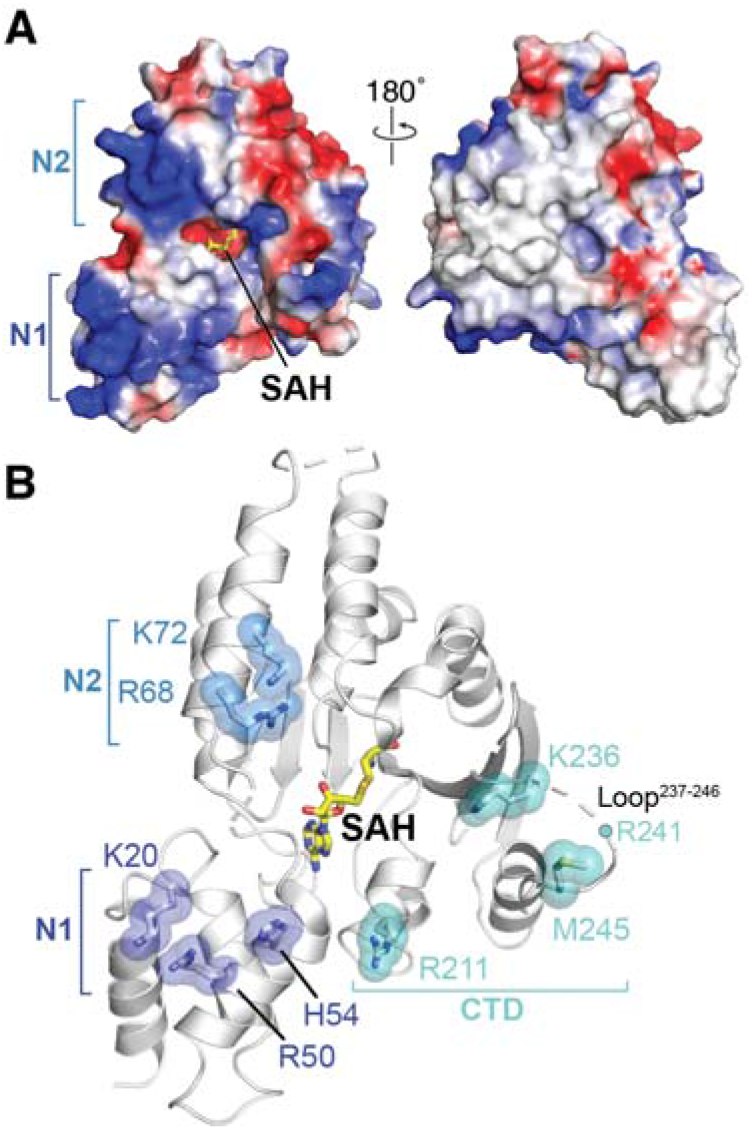
Selection of mutants defining the 30S interaction surface. **A.** The electrostatic surface potential of the RmtC structure reveals the N1 and N2 domains to be rich in positively charged residues (blue). **B.** Locations of positively charged residues in the N1 and N2 domain and other conserved or putative functionally critical residues for 30S interaction. All residues shown as sticks with semi-transparent spheres, as well as R241 located in the partially disordered Loop^237-246^) were substituted to test their role in 30S recognition (see main text and **Tables 2** and **3** for details).

To gain deeper insight into 30S recognition by RmtC and other m^7^G1405 methyltransferases, we therefore selected nine individual residues for site-directed mutagenesis, based on insights from both our RmtC structure and the previous studies of other enzymes (**Table 2** and **Fig. 3*B***). Four basic residues in the N1 and N2 domains (Lys20, Arg50, Arg68 and Lys72) were substituted with Glu to assess the contribution of the positive surface they collectively form (**Fig. 3*A***). Lys236 in the CTD is conserved in all intrinsic and acquired enzymes, while the remaining residues tested, His54 in the N1 domain and Arg211, Arg241 and Met245, were previously identified in RmtB (16). Finally, since Arg241 and Met245 were previously tested only as part of a variant in which the Loop^237-246^ was replaced by four Ala residues (16), we prepared each individual residue substitution as well as the equivalent loop alteration. All variant RmtC proteins were expressed and could be purified as for wild-type RmtC. As a further quality control to ensure that residue substitutions did not substantially impact protein folding and stability, the unfolding inflection temperature (T_i_) was determined for all purified proteins (see **Fig. S1** and **Table S1** in the Supporting Information). Almost all T_i_ values for both apparent unfolding transitions were < 2.5 °C different from wild-type RmtC, indicative of retained structural integrity. The only exception was for the RmtC-K20E/R50E double variant which exhibited slightly larger ΔT_i_ values (4.0 and 4.5 °C).

**Table 2.** Conservation of putative residues for 30S binding.

| Residue<br>(in RmtC) | % Conservation (residue) |  |  | Tested in | Refs. |
| --- | --- | --- | --- | --- | --- |
|  | All m<sup>7</sup>G1405<sup>a</sup> | Intrinsic | Acquired |  |  |
| K20 | 88 (K/R) | 100 (R) | 91 (K) | RmtC | This work |
| R50 | 100 (R/K) | 100 (K) | 100 (R/K) | Sgm | (19) |
| H54 | 96 (H) | 100 (H) | 100 (H) | RmtB | (16) |
| R68<sup>b</sup> | *nc* | *nc* | *nc* | RmtC | This work |
| K72<sup>b</sup> | *nc* | *nc* | *nc* | RmtC | This work |
| R211 | 40 (R)/40 (Q) | *nc* | 73 (R)/18 (Q) | RmtB | (16) |
| K236 | 72 (R/K) | 100 (K) | 100 (R/K) | RmtC | This work |
| R241 | 96 (R/K) | 100 (R) | 100 (R/K) | Sgm, RmtB | (16,18,19) |
| M245 | 96 (M) | 100 (M) | 100 (M) | RmtB | (16)<sup>c</sup> |

<sup>a</sup>Including sequences from aminoglycoside-producing bacteria (Intrinsic), pathogen-acquired enzymes (Acquired) and uncharacterized homologs in the chloroflexi.
<sup>b</sup>Residues are conserved only in within the RmtC subclade (7 of 8 sequences). “*nc*” indicates not conserved within the larger groups of sequences indicated.
<sup>c</sup>Residues previously only tested indirectly as part of a loop deletion variant in RmtB (residues 237-246).

As described in the following sections, each RmtC variant was assessed for 30S binding using the established FP assay and resistance (MIC) against kanamycin and gentamicin in bacteria expressing the enzymes (**Table 3**). Consistent expression of each RmtC variant was assessed under the culture conditions used for MIC measurements by immunoblotting using an anti-6×His antibody (**Fig. S2**). Thus, differences in resistance conferred and 30S binding affinity can be directly used to ascertain the role of each substituted residue in RmtC activity.

**Table 3.** RmtC variant activity.

| RmtC | Antibiotic MIC ($\mu\text{g/ml}$) | | 30S binding,<br>$K_i$ (nM)<sup>a</sup> |
| --- | --- | --- | --- |
|  | Kanamycin | Gentamicin |  |
| Wild-type | > 1024 | 1024 | 89.5 [72, 112] |
| K20E | < 2 | < 2 | NB |
| R50E | < 2 | < 2 | 977 [651, 1497] |
| K20E/R50E | < 2 | < 2 | NB |
| H54A | < 2 | < 2 | 75 [28, 203] |
| H54E | < 2 | < 2 | 90 [47, 169] |
| R68E | 256-512 | <2 | 1163 [545, 2969] |
| K72E | 256-1024 | 64-256 | 469 [225, 1005] |
| R68E/K72E | < 2 | < 2 | NB |
| R211A | 1024 | 128 | 75 [28, 196] |
| R211E | 4 | < 2 | 62 [21, 188] |
| K236A | > 1024 | 256-512 | 85 [47, 156] |
| K236E | 8 | < 2 | 76 [40, 146] |
| R241A | 1024 | 128 | 104 [79, 137] |
| R241E | 8 | < 2 | 99 [39, 252] |
| M245A | < 2 | < 2 | 55 [31, 99] |
| Loop<sup>237-246</sup>-A<sub>4</sub> | < 2 | < 2 | 63 [35, 114] |

<sup>a</sup>Values in parenthesis are 95% CI for the fit $K_i$ after averaging replicate measurements in each experiment (see **Table S2** for further details of $K_i$ determination). NB indicates “no binding”.

### Residues in N1 and N2 primarily contribute to RmtC-30S binding affinity

Single substitutions with Glu of each basic residue in either the N1 (K20E and R50E) or N2 (R68E or K72E) domain reduces 30S binding affinity of the protein in FP assays (**Fig. 4*A,B*** and **Table 3**). The extent of the reduction in binding affinities range from ~5-fold for K72E to ~11-13-fold for R50E and R68E, while no binding was measurable for K20E. Consistent with these observations, double substitutions of each pair of residues in N1 or N2 also resulted in affinities below the detectable limit in the assay (**Fig. 4*A,B*** and **Table 3**), revealing the collective contributions of the two N2 residues (Arg68 and Lys72) to binding in addition to the N1 domain (Lys20 and Arg50).

**Fig. 4.**
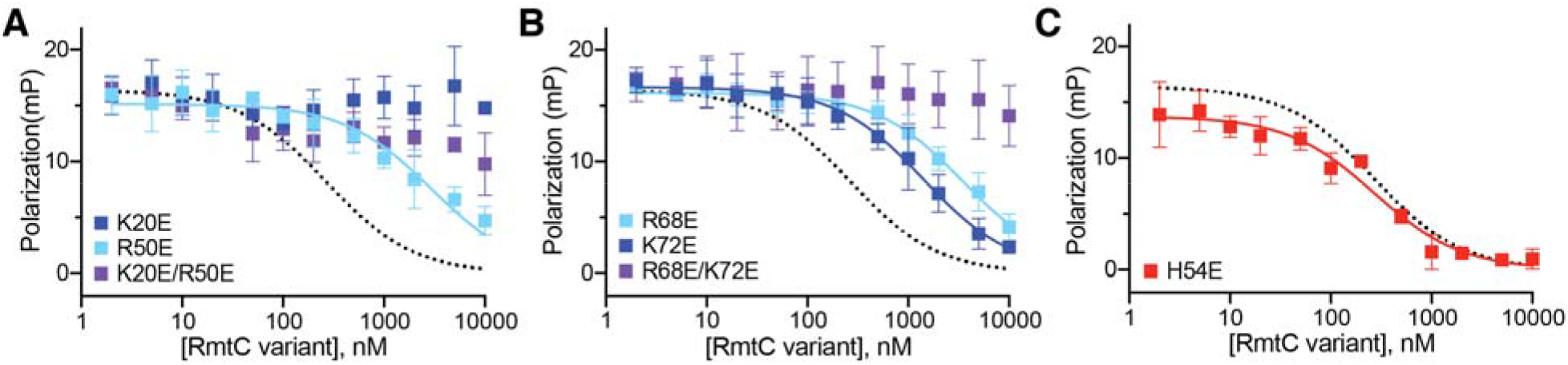
Functionally critical residues in the NTD contribute primarily to 30S binding affinity. Competition FP binding experiments with unlabeled RmtC proteins with single or double substitutions of basic (Arg/ Lys) residues with Glu in the **A.** N1 subdomain (K20E and R50E) or **B.** N2 subdomain (R68E and K72E). **C.** Competition FP binding experiment with RmtC-H54E. In all panels, the wild-type RmtC fit shown for comparison (dotted black line) is the same as that shown in **Fig 1*C,D***. Error bars represent the SEM. Binding affinity (Ki) for each variant protein derived from these data are shown in **Table 3**.

The RmtC proteins with N2 substitutions R68E, K72E or R68E/K72E were next tested for their ability to confer resistance to kanamycin or gentamicin. Intermediate MICs were determined for the single substitutions indicating a partial loss of conferred resistance, while resistance was completely abolished in the double variant (**Table 3**). The activities of these RmtC variants in bacteria thus correlated well with the measured *in vitro* binding affinities. The effects of substitutions in the N1 domain were also largely consistent in their impact on binding and activity (MIC), though it is noteworthy that the R50E substitution completely restored susceptibility to both antibiotics despite only partially reducing the enzyme’s 30S affinity (**Table 3**). This distinction may reflect a more complex role for Arg50 involving both a contribution to 30S binding affinity and a functionally critical conformational change in enzyme or substrate. For example, Arg50 might promote or stabilize a movement of the N1 subdomain relative to the CTD, as suggested by structural comparisons between RmtC and other enzymes (as noted above).

Finally, among the N1 subdomain variants, substitutions at His54 (to either Ala or Glu) produce the most striking results. For both variants, the enzyme is completely inactive, with MICs for both antibiotics at the same level as in the absence of enzyme, and yet neither substitution impacts 30S binding affinity (**Fig. 4*C*** and **Table 3**). Thus, while clearly critical for RmtC activity, H54 does not directly contribute to 30S binding, but instead must play a distinct, critical role within the substrate recognition mechanism. This observation, along with the impacts of K20E and R50E, also further points to the primary importance of the N1 subdomain in specific 30S recognition.

### Conserved CTD residues surrounding the SAM-binding pocket are functionally critical but do not contribute to 30S binding affinity

The RmtC CTD contains several residues and a structurally disordered loop region (Loop^237-246^) that are potentially critical for 30S binding. These residues line the protein surface adjacent to His 54 of the N1 domain and surrounding the opening to the SAM-binding pocket (**Table 2** and **Fig. 3*B***). Consistent with prior analyses of RmtB (16), replacement of the RmtC loop with four Ala residues (Loop^237-246^→A_4_) ablated the enzyme’s ability to confer resistance to kanamycin and gentamicin, with the same result also observed for the single substitution M245A within the loop (**Table 3**). Single substitutions to either Ala to Glu were also made for three basic residues: one within Loop^237-246^ (Arg241), one immediately preceding the loop (Lys236) and a third more distant in primary sequence but on the adjacent protein surface (Arg211). Each substitution had the same impact on protein activity in all three cases. Substitution with Ala resulted in a partial reduction in resistance conferred by the RmtC variant to kanamycin and/ or gentamicin (intermediate MICs), while substitution with Glu fully ablated resistance for all three variant enzymes (**Table 3**).

These results confirm the functional importance of the four tested residues, which line a continuous surface with H54 and the other critical residues of the N1 domain (**Fig. 3*B***). The relative effects of Ala and Glu substitutions for each of the three basic residues, R2111, K236 and R241, further suggest direct contact with the negative phosphate backbone of 16S rRNA given the greater defect with the charge reversal. Remarkably, however, none of the substitutions nor the loop swap (Loop^237-246^→A_4_) resulted in a measurable change in 30S binding affinity (**Fig. 5** and **Table 3**). Thus, like the N1 residue His54, these residues do not directly contribute to 30S binding affinity, and instead must play a distinct but critical role in substrate recognition, such as promoting or stabilizing a conformationally altered state of the enzyme and/ or substrate necessary for catalysis of m^7^G1405 modification.

**Fig. 5.**
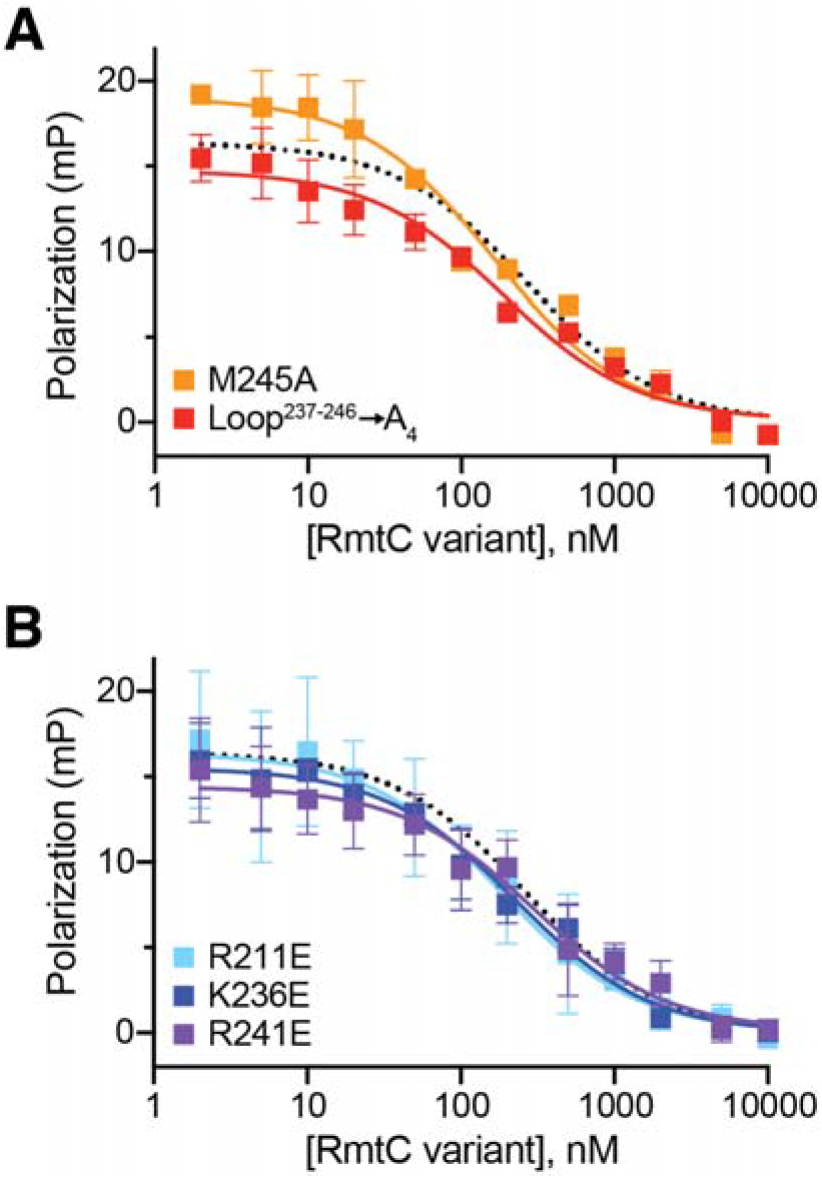
Functionally critical CTD residues do not contribute to 30S binding affinity. Competition FP binding experiments with unlabeled RmtC CTD variant proteins. **A.** Analysis of RmtC with Loop^237-246^ →A_4_ (red) or M245A single substitution with the loop. **B.** Analysis of RmtC proteins with single substitutions of basic (Arg/ Lys) residues with Glu within the CTD. In both panels, the wild-type RmtC fit shown for comparison (dotted black line) is the same as that shown in **Fig 1*C,D***. Error bars represent the SEM. Binding affinity (K_i_) for each variant protein derived from these data are shown in **Table 3**.

To gain direct insight into whether RmtC and other enzymes of this family disrupt the 30S structure upon binding, we screened a number of 30S-Rmt complexes for their suitability for single-particle cryo-electron microscopy (cryo-EM) analysis. Although strong preferred particle orientation currently precludes high-resolution 3D reconstruction, 2D class averages generated from images of a 30S-RmtG complex stabilized using the SAM analog sinefungin, clearly show disordering of the subunit head domain in the presence of the methyltransferase (**Fig. 6*A***). Thus, consistent with our interpretation of the biochemical analysis described above, m^7^G405 methyltransferase binding near the top of h44 causes significant disruption of the surrounding 16S rRNA structure, presumably allowing access to the relatively buried G1405 nucleotide for modification (**Fig. 6*B***).

**Fig. 6.**
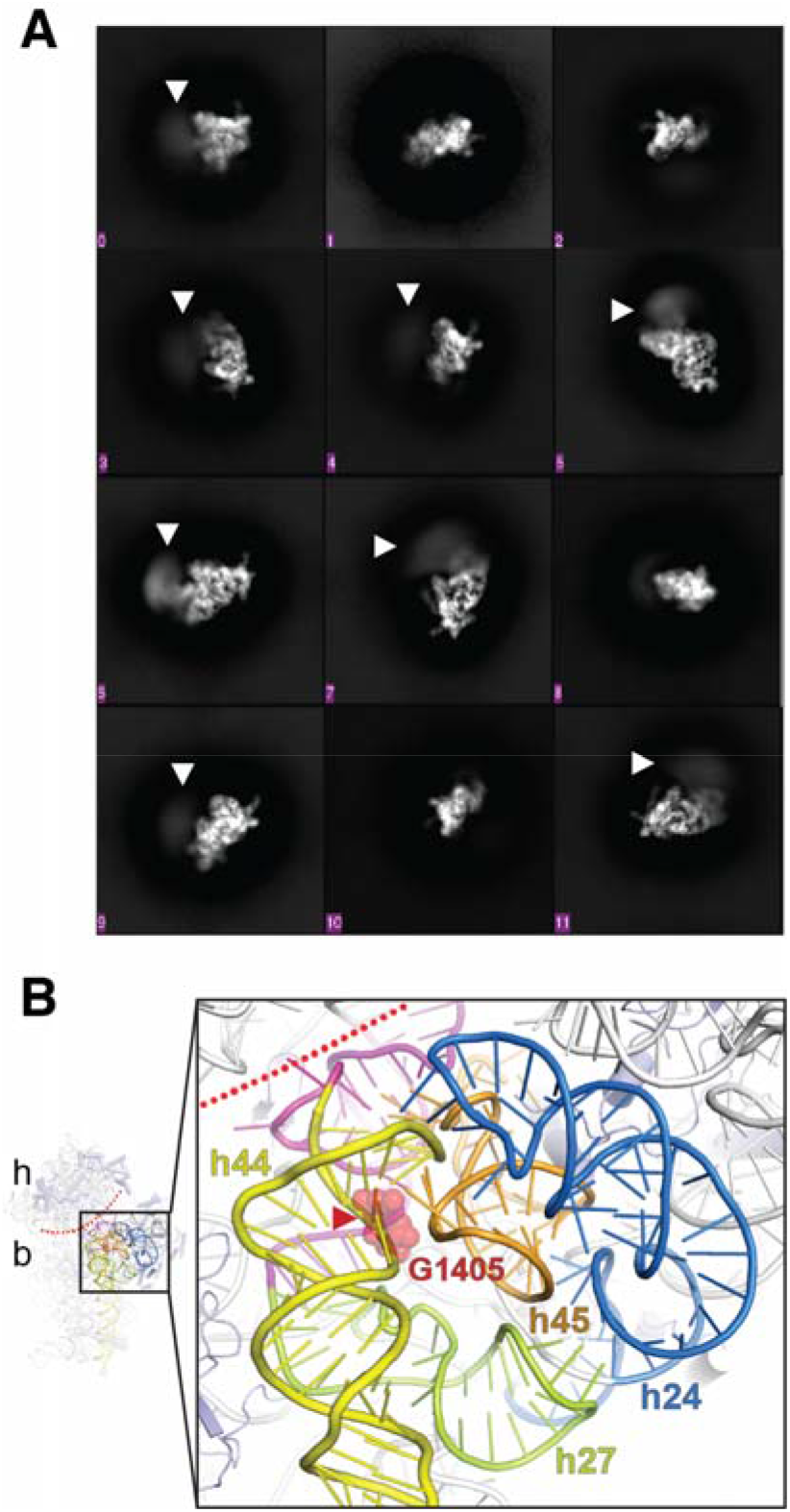
Cryo-EM analysis of a 30S-Rmt complex. ***A***, 2D class averages showing different orientations of the 30S subunit. Disorder (blurring) of the 30S head readily apparent in multiple averages (white arrowheads). ***B***, Zoomed view of the proposed 30S methyltransferase binding site architecture (generated using *E. coli* 30S, PDB ID 4V4Q). The conserved 16S rRNA tertiary surface formed by helices h24 (blue), h27 (green) and h45 (orange) is adjacent to h44 (yellow) containing the G1405 target nucleotide (red spheres). The buried location of the modified N7 atom is marked with a red arrowhead and the approximate boundary between the 30S head and body is marked with a dotted red line. Nucleotides following h27 and preceding h44, which base pair at the head-body boundary, are colored in magenta.

## DISCUSSION

The bacterial ribosome is a major target for antibiotics such as aminoglycosides, which typically interfere with the fidelity of mRNA decoding (20, 21). Although side effects have limited aminoglycoside use to treatment of serious infections, increasing resistance to other widely used antibiotics has led to a reevaluation of their use in clinical practice (21–23). Additionally, progress in mitochondrial ribosome structural biology (24, 25) and semi-synthesis of novel aminoglycosides (26, 27) can support future efforts to design new aminoglycosides with fewer side effects. As such, this important class of antimicrobials has the potential to be exceptionally useful in the treatment of serious hospital-based infections, especially those caused by Gram negative pathogens. Unfortunately, however, the clinical emergence over the last decade of aminoglycoside-resistance 16S rRNA (m^7^G1405) methyltransferases (ArmA or RmtA-H) (2, 5) pose a new threat to the efficacy of both current and new aminoglycosides, such as plazomicin (6, 7). Detailed studies, such as those described here, of the resistance methyltransferases that incorporate these rRNA modification are thus needed to support development of strategies to counter the effects of these resistance determinants.

Previous studies of m^7^G1405 methyltransferases of pathogenic (RmtB) or aminoglycoside-producer (Sgm) bacterial origin, have begun to reveal some details of 30S substrate recognition by this enzyme family (16–19). However, prior studies have typically relied on enzyme activity (e.g. MIC) measurements to indirectly infer the importance of specific residues in 30S binding. Without direct analysis of specific contributions of key residues to 30S binding affinity or other distinct roles in the process of specific substrate recognition, our understanding of the mechanism of 30S recognition and modification by the m^7^G1405 methyltransferases remained incomplete. We therefore adapted a previously developed FP assay (13) and used it here to more fully define substrate recognition by the m^7^G1405 methyltransferase enzymes.

The applicability of our FP assay using a probe based on the m^1^A1408 methyltransferase NpmA to the analysis of m^7^G1405 methyltransferase-30S interaction clearly demonstrates that the 30S binding site of these two groups of enzymes must substantially overlap. Both the m^1^A1408 and m^7^G1405 methyltransferases require the intact 30S subunit as their minimal substrate and the molecular basis for this requirement was revealed for the former enzyme subfamily by the structure of the 30S-NpmA complex. NpmA interacts exclusively with 16S rRNA and docks onto a conserved rRNA tertiary surface comprising helices 24, 27 and 45, adjacent to the h44 target site (10). This surface is bound by a group of positively charged residues, Lys66, Lys67, Lys70 and Lys71, that line a single helical region on the β2/β3 linker of the core methyltransferase fold (13). Our results with RmtC suggest that the m^7^G1405 methyltransferases likely exploit the same conserved rRNA tertiary surface for specific substrate recognition and that this is likely accomplished via interactions made by residues of the N1 and N2 domains. Specifically, a group of basic residues, Lys20, Arg50, Arg68 and Lys72, form a single positively charged surface and each contributes directly to 30S binding affinity. Lys20 and Arg50 in the N1 subdomain are highly conserved in all m^7^G1405 methyltransferases further underscoring their importance in 30S binding. In contrast, Arg68 and Lys72 in the first α-helix of the N2 domain are conserved only within the subclade comprising RmtC enzymes. However, in other m^7^G1405 methyltransferases, alternative basic residues positioned on the same surface of the protein may provide equivalent interactions with 16S rRNA, such as Lys76/Lys85 of the second α-helix of the N2 domain RmtB or Arg97/Arg106 of the second and third α-helices of the N2 domain Sgm. Thus, while some specific details may vary among different representatives of the m^7^G1405 methyltransferase subfamily, the extended positive surface created by residues of the N1/N2 domain is likely a critical first step in enzyme-substrate interaction.

Our results also reveal that multiple residues on the adjacent protein surface that surrounds the SAM-binding pocket, including His54 of the N1 subdomain and several others in the CTD, do not contribute to 30S binding affinity despite being critical for RmtC activity. These residues play no obvious direct role in RmtC protein structure and do not interact with SAM; in fact, despite their functional importance, Lys236, Arg241 and Met245 are in or adjacent to Loop^237-246^ which is disordered in the free protein. These observations and our findings that alteration of these residues abrogates activity but has no effect on 30S binding affinity suggest that they must play a distinct but essential role in substrate recognition. In NpmA, a single residue, Arg207, exhibits similar properties. Despite making no contribution to 30S binding affinity, Arg207 is nonetheless critical as it directly stabilizes the rRNA backbone of the flipped A1408 nucleotide. Our results suggest similar roles for these conserved residues in RmtC. The three basic residues, Arg211, Lys236, Arg241, likely contact the 16S rRNA backbone to stabilize a binding induced change in its structure. These residues and His54 and Met245 may also directly contact the G1405 nucleotide to position it for catalysis of methyltransfer.

Why multiple residues are required in this enzyme family compared to the single residue used by NpmA is unclear. However, our initial evidence from 2D cryo-EM class averages suggests that distortion of the 16S rRNA is large enough to cause disorder of the 30S head and body. It is noteworthy that G1405 is much less directly accessible at the top of h44 than A1408 and may thus require greater distortion of h44 and the surrounding 16S rRNA structure to create a conformation compatible with G1405 methylation by the enzyme. Reducing head-body interactions as we observe would likely allow opening of h44 near the target site and make G1405 accessible for modification, most likely via “base flipping” into the enzyme active site. We also note that nucleotides following h27, which is part of the conserved tertiary 16S rRNA surface recognized by these enzymes, are buried behind h44 near G1405 and extend to the 30S head-body boundary where they interact with residues that precede h44 (**Fig. 6*B***). Thus, a plausible mechanism is that binding to this region could relay distortion of the 16S rRNA to h44 and the 30S head-body interface. The need for this major reorganization of 16S rRNA for G1405 methylation also explains why previous attempts to dock Sgm on the 30S subunit resulted in no models with the target base within 15 Å of the SAM methyl group (17).

In summary, our model for m^7^G1405 methyltransferase action on the 30S parallels that previously developed for the m^1^A1408 methyltransferase NpmA: initial binding to 30S is mediated by multiple residues of the N1 and N2 subdomains (analogous to the NpmA β2/β3 linker), and an extended surface adjacent to this docking point is necessary to promote and/ or stabilize a novel, binding-induced 16S rRNA conformation. In particular, we speculate that one or more of the functionally critical residues on this surface is likely essential for stabilizing G1405 in a flipped conformation for methylation, as commonly observed for other RNA modifying enzymes (10, 28–30). A high-resolution structure of a 30S:m^7^G1405 methyltransferase will be necessary to define these specific molecular details. However, our findings suggest that multiple aspects of m^7^G1405 methyltransferase-substrate binding and specific recognition will emerge that may present suitable molecular targets to interfere with the action of these resistance determinants in pathogenic bacteria.

## EXPERIMENTAL PROCEDURES

### Sequence analysis

m^7^G1405 methyltransferases sequences were retrieved by BLAST search using RmtB (UniProt ID: Q76G15) as the query sequence. Sequence redundancy was removed using CD-HIT (31) with a cut off of 98% sequence identity and aligned using CLUSTAL omega. A neighbor joining phylogenetic tree was constructed using MEGA 6.0 (32) and the residue propensities were calculated using BioEdit (33).

### Protein expression and purification

Constructs for expression of RmtA (Uniprot ID: Q8GRA1), RmtB and RmtC (Uniprot ID: Q33DX5) from a modified pET44 plasmid (“pET44-HT”) were generated using synthetic *E. coli* codon-optimized genes (GenScript) as described previously (34). Equivalent expression constructs for RmtD (Uniprot ID: B0F9V0) and RmtD2 (Uniprot ID: A0A0U3JA93) were previously reported (35). Variants of RmtC were prepared using the megaprimer whole-plasmid PCR method (13, 36) and confirmed by automated DNA sequencing. Expression of all wild-type methyltransferases and variant RmtC proteins from the modified pET44 vector produced proteins with an N-terminal 6xHis tag and thrombin protease recognition sequence. For all experiments other than structural studies, proteins were used directly as the presence of the N-terminal sequence did not affect methyltransferase activity. For crystallization of RmtC, a construct for expression of tag-free wild-type RmtC (pET44-RmtC) was also generated, essentially as described previously (34).

Recombinant proteins were expressed and purified to near homogeneity using Ni^2+^-affinity and gel filtration chromatographies, as described previously (35). Purified proteins were concentrated to ~1 mg/ml, and flash frozen for storage at −80 °C before use. Tag-free wild-type RmtC was expressed similarly except that terrific broth was used as the bacterial growth medium. Purification was accomplished using heparin-affinity chromatography in 20 mM HEPES buffer pH 7.6, containing 150 mM NaCl, 10% glycerol 6 mM β-mercaptoethanol. After washing with eight column volumes of buffer, the protein was eluted using a 0.15-1 M NaCl gradient in the same buffer. Fractions containing RmtC were pooled, concentrated and the protein further purified by gel filtration chromatography on a Superdex 75 16/60 gel filtration column preequilibrated with the same buffer but containing no glycerol. Tag-free wild-type RmtC was stored as noted above or used directly for crystallization experiments (see below).

### T_i_ measurements

The thermal stability of wild-type and variant RmtC proteins was assessed using a Tycho NT.6 instrument (NanoTemper) to ensure protein quality between different preparations of proteins and before/ after freezing. In this assay, protein unfolding over a 35 to 95 °C temperature ramp is monitored via intrinsic fluorescence at 350 and 330 nm and the “inflection temperature” (T_i_) determined for each apparent unfolding transition from the temperature-dependent change in the ratio of these fluorescence measurements. All RmtC proteins unfolded in two similar apparent transitions (**Fig. S1**); T_i_ values reported in **Table S1** are the average of two measurements and replicates were typically the same within 0.5 °C.

### FP assay (K_i_ determination)

Preparation of 30S ribosomal subunits from *E. coli* (MRE600), generation of the fluorescein-labeled NpmA probe (NpmA*) and measurement of 30S-Rmt binding were accomplished essentially as previously described (13). Briefly, FP measurements were made using a Biotek Synergy Neo2 instrument with each 100 μl binding reaction containing 30S (50 nM), NpmA* (50 nM) and Rmt protein (2 nM-10 μM) in 20 mM HEPES buffer, pH 7.0, containing 75 mM KCl, 5 mM Mg(OAc)_2_, 2 mM NH_4_Cl and 3 mM β-mercaptoethanol. Solutions containing 30S and NpmA* were mixed first, incubated for 10 minutes at room temperature, aliquoted into the 96 well plate and FP measured to ensure uniform and stable FP signal prior to addition of the competing protein and final FP measurement. Initial experiments indicated equilibration was reached quickly after the addition of competing protein and was stable for at least 20 minutes. Therefore, for subsequent assays, FP was measured immediately and then each minute over the following ~5-minute period of incubation at 25 °C to ensure the system was at equilibrium with no major variations in the readings. These replicate readings were then averaged. Data handling, curve fitting to determine K_i_ values and error calculations were performed in GraphPad Prism8. All binding measurements were made in at least three independent assays for wild-type Rmt enzymes and at least two independent assays for all RmtC variants. Each assay comprised three or four replicate experiments which were separately prepared and measured but used the same preparations of protein, 30S, etc. These replicates were averaged prior to fitting in GraphPad Prism to yield the K_i_ values reported in **Table 3**. K_i_ values from fits performed on the individual values from each independent experiment were within ~2-fold agreement or better for all variants except K236A (**Table S2**). Data were fit in GraphPad Prism 8 using the “one site-fit K_i_” competition binding model:

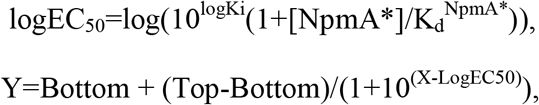

where [NpmA*] and K_d_^NpmA*^ are the concentration (in nM) and equilibrium dissociation of the labeled probe (NpmA*). Control experiments with the established competitor NpmA (13) or wild-type RmtC were included in all experiments to measure binding of the different Rmt enzymes and RmtC variants, respectively.

### Crystallization, X-ray data collection and structural refinement of the RmtC-SAH complex

Tag-free wild-type RmtC was concentrated to 12 mg/ml in the final purification buffer and mixed with a two-fold molar excess of SAH for 10 minutes at room temperature prior to screening for crystallization conditions on a Crystal Phoenix (Art Robbins Instruments). Initial crystals were obtained at 20 °C using a 1:1 mixture of protein solution and 0.1 M HEPES pH 7.0 buffer containing 2 M ammonium sulfate. An additive screen was used to further optimize crystal size and diffraction with the best diffracting crystal coming from a condition containing 3 mM mellitic acid. X-ray data were collected remotely at the Southeast Regional Collaborative Access Team (SER-CAT) 22-ID beamline at the Advanced Photon Source, Argonne National Laboratory. Data were processed and scaled using X-ray detector software (XDS) (37) in space group P61. The structure was determined by molecular replacement in Phenix (38) using a structure of *apo* RmtC (PDB ID: 6CN0) that was deposited into the PDB during the course of this study. The ligand docking and model optimization was accomplished using multiple rounds of refinement and model adjustment in Phenix (38) and Coot (39), respectively. PDB-Redo (40) was also used to optimize the quality of the final model. Complete X-ray data collection and refinement statistics are provided in **Table 1**.

### Kanamycin and gentamicin MIC assays

Fresh lysogeny broth (5 ml) containing 100 μg/ml ampicillin was inoculated (1:100 dilution) with saturated overnight culture of *E. coli* BL21(DE3) harboring pET-HT plasmid encoding wild-type or variant RmtC. Cells were grown to OD_600_ ~0.1 at 37 °C with vigorous shaking. Cells from 1 ml of this culture were collected by centrifugation, washed twice with phosphate buffered saline solution (0.5 ml) and resuspended in cation-adjusted Mueller-Hinton (CA-MHB) medium to an OD_600_ of 0.1 (5 x 10^7^ cfu/ml). Cells were further diluted 50-fold with CA-MHB and 100 μl used to inoculate (1 x 10^5^ cfu/ well) an equal volume of CA-MHB media containing 10 μM IPTG and 4-2048 μg/ml antibiotic that was pre-dispensed on a 96 well plate. For each RmtC protein, four to six individual colonies were tested from at least two independent transformations of bacterial cells with plasmid. Wells with no antibiotic or no cells served as controls in each replicate. Plates were incubated at 37 °C with shaking and OD_600_ measurements taken after 24 hours. The MIC was defined as the lowest concentration of antibiotic that inhibited growth (OD600 of <0.05 above background).

To ensure all variant proteins were comparably expressed in the MIC assay, cultures were grown on microplates under identical conditions but without antibiotic. After confirming all cultures had similar final cell densities (OD_600_ ~0.45 to 0.5) at 24 hours growth, pelleted cells were resuspended in 100 μl of 2×SDS loading dye and 5 μl loaded per lane after boiling to lyse cells and denature the proteins. His-tagged RmtC proteins were detected by immunoblotting with a rabbit anti-6×His antibody (α6×His; Proteintech; 10001-0-AP) overnight at 4 °C. Blots were probed for 1 hour at room temperature with a horseradish peroxidase-conjugated goat anti-rabbit secondary antibody (Sigma-Aldrich; A0545) treated with enhanced chemiluminescence reagent (Thermo Fisher) and imaged on a Bio-Rad ChemiDoc™ imager.

### Cryo-EM

30S-RmtG complex was crosslinked by addition of 25 μl glutaraldehyde (0.04%) and incubated for 20 minutes on ice before quenching by addition of 10 μl of glycine (0.16 mM), pH 7.4. Sample (3.5 μl) was applied to glow discharged holey-carbon quantifoil grids (R 2/2, Cu 200) and blotted for 3.5 s at 95% humidity and 8 °C before vitrification by plunging into liquid ethane using a FEI Vitrobot Mark IV. A FEI Titan Krios microscope operating at 300 kV and equipped with a K3 direct detector camera (Gatan) was used to collect 1191 movie frames with Leginon (41). Sixty frames per movie were collected at the total dose of 60.13 e^−^/Å^2^ on the sample. The magnification was 81,000× and the super-resolution frames were binned 2× corresponding to a pixel size of 1.11 Å.

All pre-processing steps were performed in Appion (42). MotionCor2 (43) was used to align the frames of each micrograph, correct for global and local (5×5 patches) beam-induced motion and to dose weight individual frames. Defocus values were determined using CTFFIND4 (44). An initial 3,000 particles were picked from using the reference-free particle picker DoG (45) and subjected to 2D classification in cryoSPARC (46). To generate an initial template, e2proc2d.py of EMAN2 (47) was used to compute the rotational average of the 10 best resolved classes. The resulting template was low pass filtered to 15 Å and used to extract particles from the entire set of frame-aligned micrographs using the FindEM template picking software (48). Mask diameter used for template picking was 300 Å. A total of 215,554 particles were extracted with 432×432 pixel boxes from full-size images, and the stack and metadata file were exported to cisTEM (49) and used to generate the 2D classes shown in **Fig. 6**.

## Acknowledgements

We thank members of the Conn and Dunham labs at Emory University for helpful discussions throughout the course of this work.

## FOOTNOTES

This work was supported by National Institutes of Health grants R01-AI088025 (to GLC) and T32-GM008367, and postdoctoral fellowship DEY18F0 from Cystic Fibrosis Foundation (to DD) SER-CAT is supported by its member institutions and equipment grants (S10 RR25528 and S10 RR028976) from the National Institutes of Health. Use of the Advanced Photon Source was supported by the U. S. Department of Energy, Office of Science, Office of Basic Energy Sciences, under Contract No. W-31-109-Eng-38.

## Abbreviations used are

AME: aminoglycoside modifying enzyme
CA-MHB: cation-adjusted Mueller-Hinton broth
CTD: carboxy-terminal domain
FP: fluorescence polarization
h44: (16S rRNA) helix 44
MIC: minimum inhibitory concentration
NTD: amino-terminal domain
Rmt: (aminoglycoside) resistance methyltransferase
SAH: S-adenosylhomocysteine
SAM: S-adenosyl-L-methionine
T_i_: inflection temperature

## REFERENCES

1. Cundliffe, E. (1989) How antibiotic-producing organisms avoid suicide. Annual Review of Microbiology 43, 207–233

2. Wachino, J., and Arakawa, Y. (2012) Exogenously acquired 16S rRNA methyltransferases found in aminoglycoside-resistant pathogenic Gram-negative bacteria: an update. Drug Resist Updat 15, 133–148

3. Rahman, M., Shukla, S. K., Prasad, K. N., Ovejero, C. M., Pati, B. K., Tripathi, A., Singh, A., Srivastava, A. K., and Gonzalez-Zorn, B. (2014) Prevalence and molecular characterisation of New Delhi metallo-beta-lactamases NDM-1, NDM-5, NDM-6 and NDM-7 in multidrug-resistant Enterobacteriaceae from India. Int J Antimicrob Agents 44, 30–37

4. Long, H., Feng, Y., Ma, K., Liu, L., McNally, A., and Zong, Z. (2019) The co-transfer of plasmid-borne colistin-resistant genes mcr-1 and mcr-3.5, the carbapenemase gene blaNDM-5 and the 16S methylase gene rmtB from Escherichia coli. Sci Rep 9, 696

5. Doi, Y., Wachino, J. I., and Arakawa, Y. (2016) Aminoglycoside Resistance: The Emergence of Acquired 16S Ribosomal RNA Methyltransferases. Infect Dis Clin North Am 30, 523–537

6. Aggen, J. B., Armstrong, E. S., Goldblum, A. A., Dozzo, P., Linsell, M. S., Gliedt, M. J., Hildebrandt, D. J., Feeney, L. A., Kubo, A., Matias, R. D., Lopez, S., Gomez, M., Wlasichuk, K. B., Diokno, R., Miller, G. H., and Moser, H. E. (2010) Synthesis and spectrum of the neoglycoside ACHN-490. Antimicrob Agents Chemother 54, 4636–4642

7. Livermore, D. M., Mushtaq, S., Warner, M., Zhang, J. C., Maharjan, S., Doumith, M., and Woodford, N. (2011) Activity of aminoglycosides, including ACHN-490, against carbapenem-resistant Enterobacteriaceae isolates. J Antimicrob Chemother 66, 48–53

8. Garneau-Tsodikova, S., and Labby, K. J. (2016) Mechanisms of Resistance to Aminoglycoside Antibiotics: Overview and Perspectives. Medchemcomm 7, 11–27

9. Wachino, J., Shibayama, K., Kurokawa, H., Kimura, K., Yamane, K., Suzuki, S., Shibata, N., Ike, Y., and Arakawa, Y. (2007) Novel plasmid-mediated 16S rRNA m1A1408 methyltransferase, NpmA, found in a clinically isolated Escherichia coli strain resistant to structurally diverse aminoglycosides. Antimicrob Agents Chemother 51, 4401–4409

10. Dunkle, J. A., Vinal, K., Desai, P. M., Zelinskaya, N., Savic, M., West, D. M., Conn, G. L., and Dunham, C. M. (2014) Molecular recognition and modification of the 30S ribosome by the aminoglycoside-resistance methyltransferase NpmA. Proc Natl Acad Sci U S A 111, 6275–6280

11. Husain, N., Obranic, S., Koscinski, L., Seetharaman, J., Babic, F., Bujnicki, J. M., Maravic-Vlahovicek, G., and Sivaraman, J. (2011) Structural basis for the methylation of A1408 in 16S rRNA by a panaminoglycoside resistance methyltransferase NpmA from a clinical isolate and analysis of the NpmA interactions with the 30S ribosomal subunit. Nucleic Acids Res 39, 1903–1918

12. Macmaster, R., Zelinskaya, N., Savic, M., Rankin, C. R., and Conn, G. L. (2010) Structural insights into the function of aminoglycoside-resistance A1408 16S rRNA methyltransferases from antibiotic-producing and human pathogenic bacteria. Nucleic Acids Res 38, 7791–7799

13. Vinal, K., and Conn, G. L. (2017) Substrate Recognition and Modification by a Pathogen-Associated Aminoglycoside Resistance 16S rRNA Methyltransferase. Antimicrob Agents Chemother 61

14. Savic, M., Sunita, S., Zelinskaya, N., Desai, P. M., Macmaster, R., Vinal, K., and Conn, G. L. (2015) 30S Subunit-dependent activation of the Sorangium cellulosum So ce56 aminoglycoside resistance-conferring 16S rRNA methyltransferase Kmr. Antimicrob Agents Chemother 59, 2807–2816

15. Witek, M. A., and Conn, G. L. (2016) Functional dichotomy in the 16S rRNA (m1A1408) methyltransferase family and control of catalytic activity via a novel tryptophan mediated loop reorganization. Nucleic Acids Res 44, 342–353

16. Schmitt, E., Galimand, M., Panvert, M., Courvalin, P., and Mechulam, Y. (2009) Structural bases for 16 S rRNA methylation catalyzed by ArmA and RmtB methyltransferases. J Mol Biol 388, 570–582

17. Husain, N., Tkaczuk, K. L., Tulsidas, S. R., Kaminska, K. H., Cubrilo, S., Maravic-Vlahovicek, G., Bujnicki, J. M., and Sivaraman, J. (2010) Structural basis for the methylation of G1405 in 16S rRNA by aminoglycoside resistance methyltransferase Sgm from an antibiotic producer: a diversity of active sites in m7G methyltransferases. Nucleic Acids Res 38, 4120–4132

18. Savic, M., Ilic-Tomic, T., Macmaster, R., Vasiljevic, B., and Conn, G. L. (2008) Critical residues for cofactor binding and catalytic activity in the aminoglycoside resistance methyltransferase Sgm. J Bacteriol 190, 5855–5861

19. Maravic Vlahovicek, G., Cubrilo, S., Tkaczuk, K. L., and Bujnicki, J. M. (2008) Modeling and experimental analyses reveal a two-domain structure and amino acids important for the activity of aminoglycoside resistance methyltransferase Sgm. Biochim Biophys Acta 1784, 582–590

20. Lin, J., Zhou, D., Steitz, T. A., Polikanov, Y. S., and Gagnon, M. G. (2018) Ribosome-Targeting Antibiotics: Modes of Action, Mechanisms of Resistance, and Implications for Drug Design. Annu Rev Biochem 87, 451–478

21. Becker, B., and Cooper, M. A. (2013) Aminoglycoside antibiotics in the 21st century. ACS Chem Biol 8, 105–115

22. Takahashi, Y., and Igarashi, M. (2017) Destination of aminoglycoside antibiotics in the ‘postantibiotic era’. J Antibiot (Tokyo)

23. Durante-Mangoni, E., Grammatikos, A., Utili, R., and Falagas, M. E. (2009) Do we still need the aminoglycosides? Int J Antimicrob Agents 33, 201–205

24. Amunts, A., Brown, A., Toots, J., Scheres, S. H. W., and Ramakrishnan, V. (2015) Ribosome. The structure of the human mitochondrial ribosome. Science 348, 95–98

25. Greber, B. J., Bieri, P., Leibundgut, M., Leitner, A., Aebersold, R., Boehringer, D., and Ban, N. (2015) Ribosome. The complete structure of the 55S mammalian mitochondrial ribosome. Science 348, 303–308

26. Sonousi, A., Sarpe, V. A., Brilkova, M., Schacht, J., Vasella, A., Bottger, E. C., and Crich, D. (2018) Effects of the 1- N-(4-Amino-2 S-hydroxybutyryl) and 6’- N-(2-Hydroxyethyl) Substituents on Ribosomal Selectivity, Cochleotoxicity, and Antibacterial Activity in the Sisomicin Class of Aminoglycoside Antibiotics. Acs Infectious Diseases 4, 1114–1120

27. Matsushita, T., Sati, G. C., Kondasinghe, N., Pirrone, M. G., Kato, T., Waduge, P., Kumar, H. S., Sanchon, A. C., Dobosz-Bartoszek, M., Shcherbakov, D., Juhas, M., Hobbie, S. N., Schrepfer, T., Chow, C. S., Polikanov, Y. S., Schacht, J., Vasella, A., Bottger, E. C., and Crich, D. (2019) Design, Multigram Synthesis, and in Vitro and in Vivo Evaluation of Propylamycin: A Semisynthetic 4,5-Deoxystreptamine Class Aminoglycoside for the Treatment of Drug-Resistant Enterobacteriaceae and Other Gram-Negative Pathogens. J Am Chem Soc 141, 5051–5061

28. Lee, T. T., Agarwalla, S., and Stroud, R. M. (2005) A unique RNA Fold in the RumA-RNA-cofactor ternary complex contributes to substrate selectivity and enzymatic function. Cell 120, 599–611

29. Alian, A., Lee, T. T., Griner, S. L., Stroud, R. M., and Finer-Moore, J. (2008) Structure of a TrmA-RNA complex: A consensus RNA fold contributes to substrate selectivity and catalysis in m5U methyltransferases. Proc Natl Acad Sci U S A 105, 6876–6881

30. Wang, C., Jia, Q., Zeng, J., Chen, R., and Xie, W. (2017) Structural insight into the methyltransfer mechanism of the bifunctional Trm5. Science Advances 3, e1700195

31. Li, W., and Godzik, A. (2006) Cd-hit: a fast program for clustering and comparing large sets of protein or nucleotide sequences. Bioinformatics 22, 1658–1659

32. Tamura, K., Stecher, G., Peterson, D., Filipski, A., and Kumar, S. (2013) MEGA6: Molecular Evolutionary Genetics Analysis version 6.0. Mol Biol Evol 30, 2725–2729

33. Hall, T., Biosciences, I., and Carlsbad, C. (2011) BioEdit: an important software for molecular biology. GERF Bull Biosci 2, 60–61

34. Witek, M. A., and Conn, G. L. (2014) Expansion of the aminoglycoside-resistance 16S rRNA (m(1)A1408) methyltransferase family: expression and functional characterization of four hypothetical enzymes of diverse bacterial origin. Biochim Biophys Acta 1844, 1648–1655

35. Correa, L. L., Witek, M. A., Zelinskaya, N., Picao, R. C., and Conn, G. L. (2016) Heterologous Expression and Functional Characterization of the Exogenously Acquired Aminoglycoside Resistance Methyltransferases RmtD, RmtD2, and RmtG. Antimicrob Agents Chemother 60, 699–702

36. Witek, M. A., Kuiper, E. G., Minten, E., Crispell, E. K., and Conn, G. L. (2017) A Novel Motif for S-Adenosyl-l-methionine Binding by the Ribosomal RNA Methyltransferase TlyA from Mycobacterium tuberculosis. J Biol Chem 292, 1977–1987

37. Kabsch, W. (2010) Xds. Acta Crystallogr D Biol Crystallogr 66, 125–132

38. Adams, P. D., Afonine, P. V., Bunkoczi, G., Chen, V. B., Davis, I. W., Echols, N., Headd, J. J., Hung, L. W., Kapral, G. J., Grosse-Kunstleve, R. W., McCoy, A. J., Moriarty, N. W., Oeffner, R., Read, R. J., Richardson, D. C., Richardson, J. S., Terwilliger, T. C., and Zwart, P. H. (2010) PHENIX: a comprehensive Python-based system for macromolecular structure solution. Acta Crystallographica. Section D: Biological Crystallography 66, 213–221

39. Emsley, P., Lohkamp, B., Scott, W. G., and Cowtan, K. (2010) Features and development of Coot. Acta Crystallogr D Biol Crystallogr 66, 486–501

40. Joosten, R. P., Long, F., Murshudov, G. N., and Perrakis, A. (2014) The PDB_REDO server for macromolecular structure model optimization. IUCrJ 1, 213–220

41. Suloway, C., Pulokas, J., Fellmann, D., Cheng, A., Guerra, F., Quispe, J., Stagg, S., Potter, C. S., and Carragher, B. (2005) Automated molecular microscopy: the new Leginon system. J Struct Biol 151, 41–60

42. Lander, G. C., Stagg, S. M., Voss, N. R., Cheng, A., Fellmann, D., Pulokas, J., Yoshioka, C., Irving, C., Mulder, A., Lau, P. W., Lyumkis, D., Potter, C. S., and Carragher, B. (2009) Appion: an integrated, database-driven pipeline to facilitate EM image processing. J Struct Biol 166, 95–102

43. Zheng, S. Q., Palovcak, E., Armache, J. P., Verba, K. A., Cheng, Y., and Agard, D. A. (2017) MotionCor2: anisotropic correction of beam-induced motion for improved cryo-electron microscopy. Nat Methods 14, 331–332

44. Rohou, A., and Grigorieff, N. (2015) CTFFIND4: Fast and accurate defocus estimation from electron micrographs. J Struct Biol 192, 216–221

45. Voss, N. R., Yoshioka, C. K., Radermacher, M., Potter, C. S., and Carragher, B. (2009) DoG Picker and TiltPicker: software tools to facilitate particle selection in single particle electron microscopy. J Struct Biol 166, 205–213

46. Punjani, A., Rubinstein, J. L., Fleet, D. J., and Brubaker, M. A. (2017) cryoSPARC: algorithms for rapid unsupervised cryo-EM structure determination. Nature Methods 14, 290

47. Tang, G., Peng, L., Baldwin, P. R., Mann, D. S., Jiang, W., Rees, I., and Ludtke, S. J. (2007) EMAN2: an extensible image processing suite for electron microscopy. J Struct Biol 157, 38–46

48. Roseman, A. M. (2004) FindEM--a fast, efficient program for automatic selection of particles from electron micrographs. J Struct Biol 145, 91–99

49. Grant, T., Rohou, A., and Grigorieff, N. (2018) cisTEM, user-friendly software for single-particle image processing. Elife 7

