## Supporting Information for "Functionally critical residues in the aminoglycoside resistance-associated methyltransferase RmtC play distinct roles in 30S substrate recognition"

Running title: *30S substrate recognition by RmtC*

**Keywords**: antibiotic resistance, ribosomal RNA (rRNA), ribosome, RNA methylation, RNA methyltransferase.

---

**Table S1.** Analysis of wild-type and variant RmtC protein folding.

**Fig. S1.** Analysis of wild-type and variant RmtC protein folding and stability using thermal unfolding.

**Fig. S2.** Expression of wild-type and variant RmtC proteins under the culture conditions used for antibiotic MIC measurements.

**Table S2.** Analysis of 30S-RmtC variant binding by competition FP.

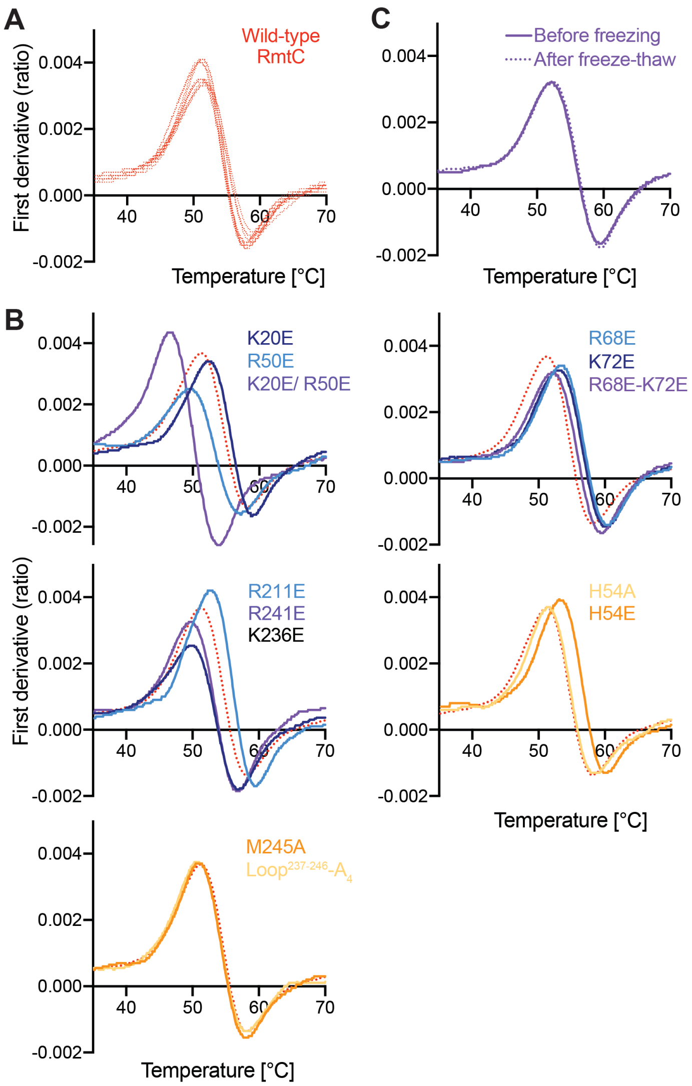


**Table S1. Analysis of wild-type and variant RmtC protein folding.**

| RmtC | Unfolding $T_i$ (°C)[a] | |
|---|---|---|
| | 1st | 2nd |
| Wild-type | 51.3 (-) | 58.0 (-) |
| K20E | 52.4 (1.1) | 59.1 (1.1) |
| R50E | 49.5 (1.8) | 57.1 (0.9) |
| K20E/R50E | 46.8 (4.5) | 54.0 (4.0) |
| H54A | 51.4 (0.1) | 58.5 (0.5) |
| H54E | 53.5 (2.2) | 60.3 (2.3) |
| R68E | 53.3 (2.0) | 60.4 (2.4) |
| K72E | 53.0 (1.7) | 60.2 (2.2) |
| R68E/K72E | 52.2 (0.9) | 59.5 (1.5) |
| R211A | 52.0 (0.7) | 59.0 (1.0) |
| R211E | 52.8 (1.5) | 59.6 (1.6) |
| K236A | 50.4 (0.9) | 57.5 (0.5) |
| K236E | 49.8 (1.5) | 56.9 (1.1) |
| R241A | 51.3 (0.0) | 58.3 (0.3) |
| R241E | 49.7 (1.6) | 56.8 (1.2) |
| M245A | 51.1 (0.2) | 58.0 (0.0) |
| $Loop^{237\text{-}246}$-$A_4$ | 50.9 (0.4) | 58.1 (0.1) |

[a]Values in parenthesis are absolute differences in $T_i$ compared to wild-type RmtC.

**Fig. S1. Quality control of purified wild-type and variant RmtC proteins by thermal denaturation.** ***A***, Replicate measurements of wild-type RmtC unfolding monitored using intrinsic fluorescence at 330 and 350 nm, illustrating the reproducibility of the method between multiple experiments and preparations of protein. First derivative plots are shown for fluorescence ratio (350/ 330 nm) from which $T_i$ values were determined corresponding to the positive (51.3 ˚C) and negative (~58.0 ˚C) peaks in the unfolding profile. ***B***, Equivalent analysis for RmtC variants as indicated. In each panel, wild-type RmtC is shown for comparison (red dotted line representing the average of all measurements in *panel A*). $T_i$ values determined from the plots in *panels A* and *B* are shown in **Table S1**. ***C***, Example of protein quality control for RmtC-R68E/K72E showing unfolding profiles of the same protein preparation before and after storage at -80 ˚C.

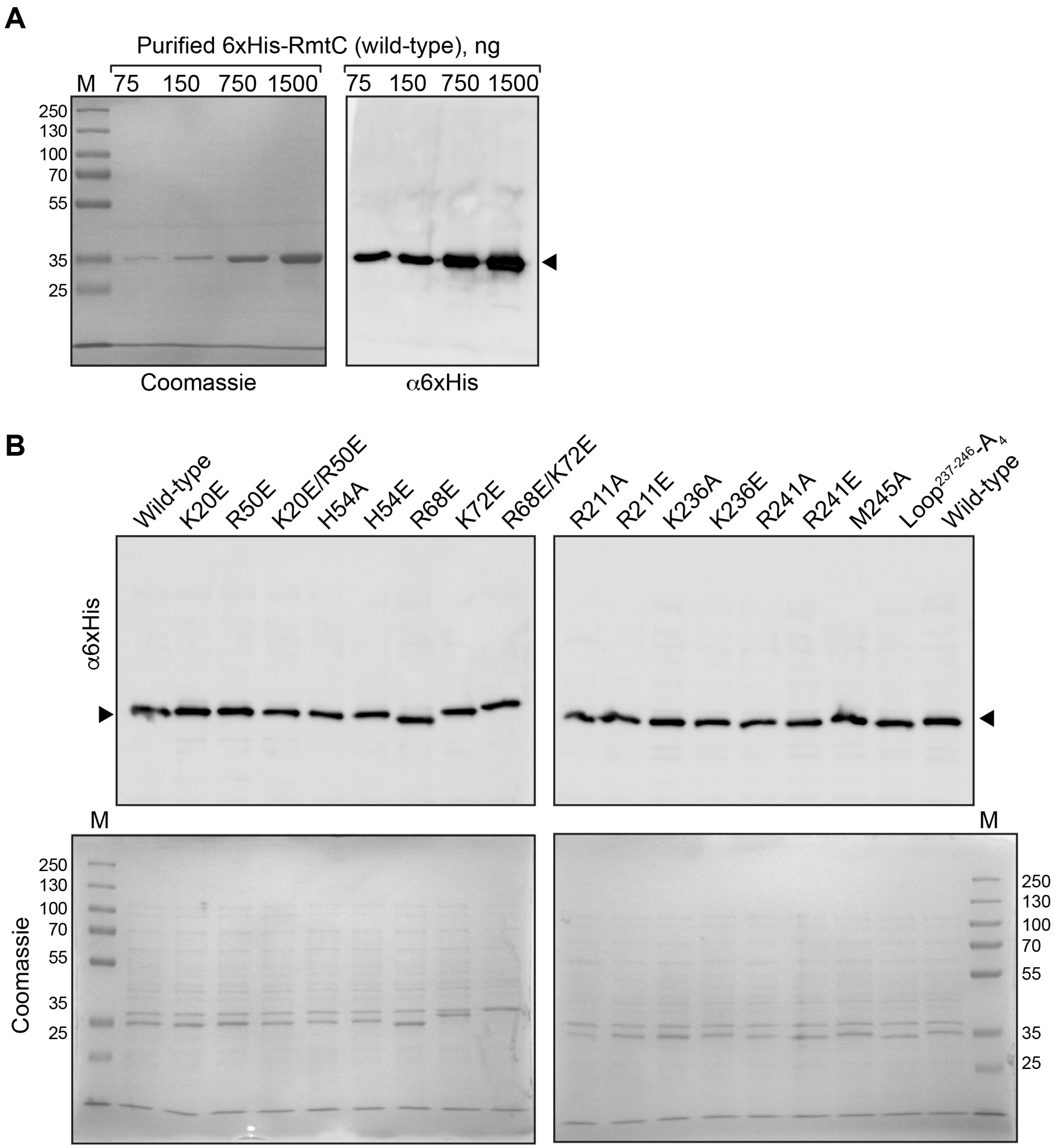


**Fig. S2. Expression of wild-type and variant RmtC proteins under the culture conditions used for antibiotic MIC measurements.** ***A***, Validation of the rabbit anti-6×His antibody (α6×His) for detection of 6×His-tagged RmtC proteins. The indicated amounts of purified wild-type 6×His-RmtC were resolved on two SDS-PAGE gels in parallel and used for staining with coomassie or immunoblotting with α6×His. ***B***, Immunoblot (top) and coomassie stained gel of the indicated RmtC proteins cultured under the conditions used for MIC assays. All variants are expressed comparably to the wild-type protein.

**Table S2. Analysis of 30S-RmtC variant binding by competition FP.**

| RmtC | Individual experiments[a] | | Both experiments (single fit)[b] | |
|---|---|---|---|---|
| | 30S binding, $K_i$ (nM)[c] | | 30S binding, $K_i$ (nM)[c] | Fit $R^2$ |
| | Expt. 1 | Expt. 2 | | |
| R50E | 913 [575, 1488] | 1015 [756, 1379] | 977 [651, 1497] | 0.86 |
| H54A | 114 [82, 157] | 61 [44, 85] | 75 [28, 203] | 0.83 |
| H54E | 82 [61, 111] | 97 [51, 182] | 90 [47, 169] | 0.85 |
| R68E | 860 [520, 1488] | 1671 [880, 3810] | 1163 [545, 2969] | 0.91 |
| K72E | 647 [426, 1000] | 349 [211, 574] | 469 [225, 1005] | 0.90 |
| R211A | 68 [38, 125] | 81 [49, 134] | 75 [28, 196] | 0.84 |
| R211E | 45 [24, 86] | 80 [54, 124] | 62 [21, 188] | 0.81 |
| K236A | 54 [36, 81] | 152 [98, 234] | 85 [47, 156] | 0.94 |
| K236E | 66 [42, 115] | 84 [54, 132] | 76 [40, 146] | 0.93 |
| R241A | 122 [86, 174] | 89 [65, 123] | 104 [79, 137] | 0.98 |
| R241E | 74 [32, 171] | 135 [87, 209] | 99 [39, 252] | 0.86 |
| M245A | 46 [18, 131] | 71 [47, 109] | 55 [31, 99] | 0.95 |
| $Loop^{237\text{-}246}$-$A_4$ | 50 [34, 73] | 93 [57, 152] | 63 [35, 114] | 0.94 |

[a]Data fit using the "One site-fit Ki" model in Graph Pad Prism 8 considering each of the 3-4 replicate measurements separately (each binding experiment was prepared independently but in parallel using the same preparations of protein, NpmA* and 30S).
[b]Data from each set of replicate measurements was averaged prior to fitting using the "One site-fit Ki" model in Graph Pad Prism 8.
[c]Values in parenthesis are 95% CI for the fit $K_i$.
